# A catalog of ancient proxies for modern genetic variants

**DOI:** 10.1101/2025.05.19.654975

**Authors:** Colin M. Brand, John A. Capra

**Affiliations:** Bakar Computational Health Sciences Institute, University of California, San Francisco, CA, USA; Department of Epidemiology and Biostatistics, University of California, San Francisco, CA, USA; Department of Bioengineering and Therapeutic Sciences, University of California San Francisco, San Francisco, CA, USA

## Abstract

The ability to observe the genomes of past human populations using ancient DNA provides an extraordinary perspective on many fundamental questions in human genetics, including understanding the evolutionary history of variants that underlie human disease and other phenotypes. However, ancient DNA is often damaged and degraded, yielding low-coverage of most nucleotides. Further, many publicly available genotypes for ancient humans are limited to ∼1.23 million specific loci. Thus, variants of interest often fall outside these specific positions, limiting the ability of ancient DNA to shed light on many loci. Here, we address this challenge by quantifying linkage disequilibrium (LD) between modern variants and ancient genotyped variants (AGVs) to generate a catalog enabling rapid identification of proxy variants. We identified 260,732,675 pairs of AGVs and modern variants with a minimum LD threshold hold of *R*^*2*^ ≥ 0.2. Even at *R*^*2*^ ≥ 0.9, ≥ 60% of common variants were linked to an AGV in non-African ancestry groups, as were 34% of common variants in Africans. We evaluated the accuracy of the genotypes inferred from proxy variants in two high-coverage ancient genomes finding that > 90% of genotypes were correctly predicted, even in a 45,000 year old individual. We also find that AGVs are significantly older than expected and that many likely are evolving neutrally. We integrate these results in a database that researchers can easily query to identify ancient proxy variants if their variant of interest is not directly genotyped in ancient humans.

## INTRODUCTION

The sequencing of ancient DNA from humans and other species offers the opportunity to directly study DNA sequences through time (Pääbo et al., 2004; Slatkin and Racimo, 2016; Willerslev and Cooper, 2004). Examining the presence/absence of alleles in different ancient populations can reveal the history of evolutionary origins and pressures on variants that underlie human disease and other phenotypes of interest (Barrie et al., 2024; Patin and Quintana-Murci, 2024). Such analyses not only shed light on recent evolutionary processes in humans but may also help uncover molecular mechanisms for selected traits (Mathieson and Mathieson, 2018; Yilmaz et al., 2025).

Recovering and sequencing ancient DNA is challenging due to degradation resulting in small fragment sizes, the presence of chemical damage, and overall low abundance of endogenous versus exogenous DNA (Orlando et al., 2021). While some ancient samples have been sequenced to coverage that is comparable to modern samples, most ancient sequences are low-coverage, often averaging less than one read per nucleotide (Orlando et al., 2021). Further, methods are often used to capture, amplify, and sequence ancient DNA from a subset of informative sites across the genome. Recently, a harmonized dataset of nearly 10,000 ancient genotypes was released: the Allen Ancient DNA Resource (AADR) (Mallick et al., 2024). This resource contains genotypes at up to ∼1.23 million loci from both ancient and modern samples. However, two barriers currently limit the utility of this resource in studying variants of interest, especially for relevant to human disease. First, the dataset is not easily searchable without expertise using population genetics command line tools. Second, variants of interest commonly fall outside these assayed positions, limiting the ability of ancient DNA to shed light on the history of many loci. Imputation of ancient DNA samples using methods developed for modern genotype data is possible in some settings (Escobar-Rodríguez and Veeramah, 2024; Garrido Marques et al., 2024; Sousa da Mota et al., 2023). However, these approaches are in early stages of development, and given the diversity of age, coverage, damage, and contamination in ancient DNA, it is unlikely that a single method will work for all samples.

Here, we addressed these challenges by building a simple searchable web tool and leveraging the fact that many modern variants occur in strong linkage disequilibrium (LD) with ancient genotyped variants (AGVs). This enabled us to generate a resource for rapid identification of whether a modern variant of interest has been genotyped in ancient samples, and if not, determining whether AGVs to use as proxy variants are available. We evaluated the number and accuracy of genotypes inferred from these variant pairs at different LD thresholds using two high-coverage ancient human genomes. Then, we quantified the age distribution of AGVs and highlight examples of different allele frequency dynamics. Finally, we traced the history of an example modern variant using an AGV proxy to understand its origin and evolution.

## RESULTS

### Identifying modern variants in linkage disequilibrium with ancient genotyped variants

Quantifying the number of modern variants that occur in LD with AGVs requires pairwise LD information from a large, diverse sample. We chose TopMed (Huang et al., 2022; Taliun et al., 2021) because of its diversity and large sample size compared to other similar datasets. This enabled the identification of linked variants that occur at low frequencies (< 0.1%) (Fig. 1A). We analyzed pairwise LD information pre-computed in the TopLD database derived from autosomal and X chromosome whole genome sequencing (WGS) data in four TOPMed cohorts: BioMe, MESA, JHS, and WHI (Huang et al., 2022; Taliun et al., 2021). Local and global ancestry were estimated per sample using reference populations from the Thousand Genomes Project—TGP (Auton et al., 2015) and the Human Genome Diversity Project—HGDP (Rosenberg et al., 2002). Global ancestry was used to categorize unrelated individuals into one of four “ancestry groups”: African, East Asian, European, and South Asian. Here, “ancestry group” refers to individuals with genetic similarity to each other and reference populations from both TGP and HGDP. LD was then estimated separately in each ancestry group for all variant pairs within 500 kb. Variant pairs with *R*^*2*^ ≥ 0.2 were retained resulting in LD information for 223,322,434 total variants (Fig. 1A).

**Fig. 1.**
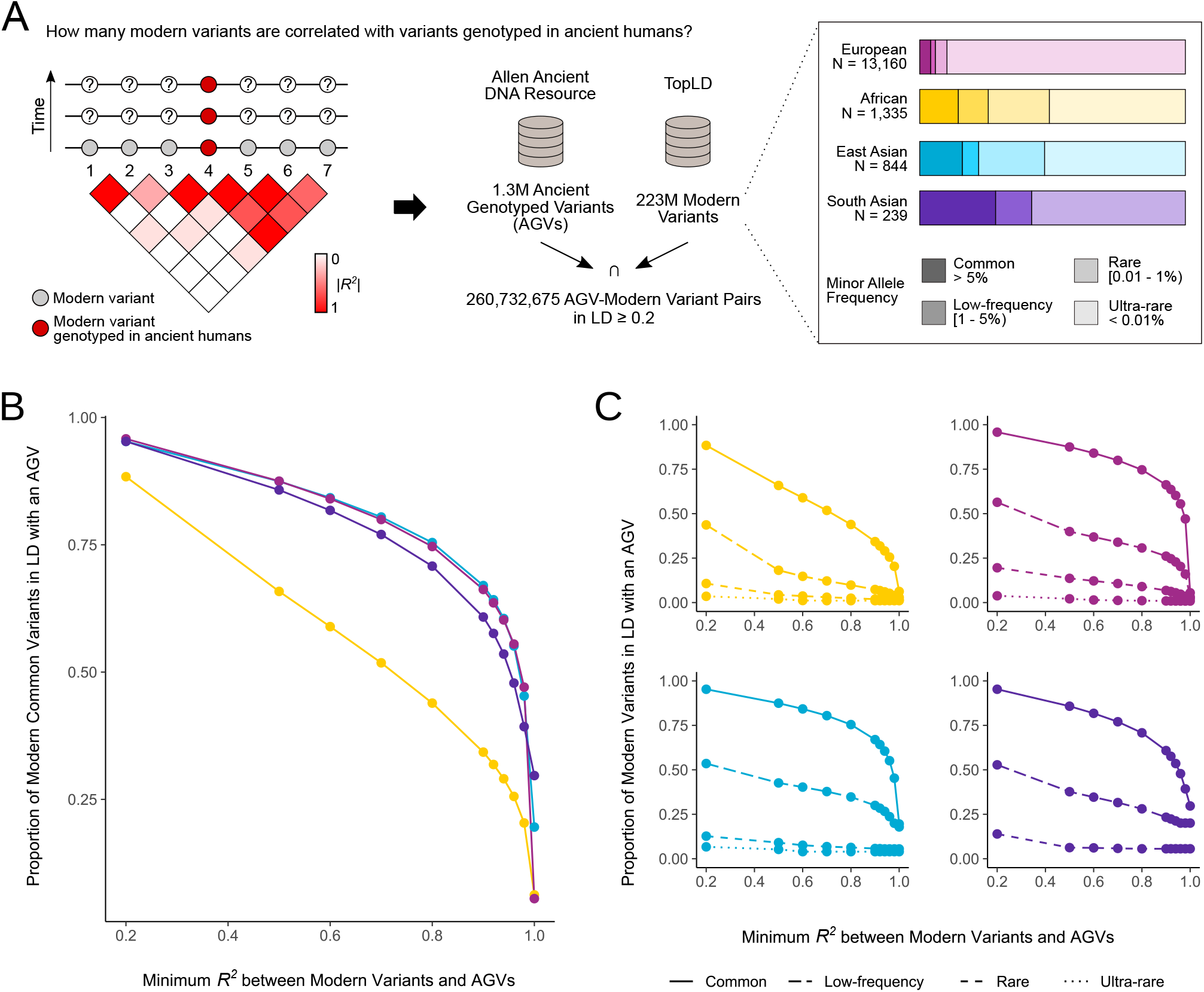
Quantifying modern variants in linkage disequilibrium with variants genotyped in ancient humans. (A) Schematic of pipeline for identifying modern variants in linkage disequilibrium (LD) with variants genotyped in ancient humans. We intersected 1,199,828 autosomal and X chromosome ancient genotyped variants (AGVs) from the Allen Ancient DNA Resource (Mallick et al., 2024) with 223,322,434 autosomal and X chromosome variants from TopLD (Huang et al., 2022; Taliun et al., 2021). We identified 260,732,675 variant pairs from TopLD where one of the variants was an AGV in at least one of four “ancestry groups”. The distribution of minor allele frequency (MAF) classes per ancestry group is quantified in the bar plot. (B) The proportion of common variants in LD with an AGV at a given threshold, stratified by ancestry group, which is indicated by color as in **A**. Proportions at LD thresholds of 0.2, 0.4, 0.6, 0.8, 0.9, 0.92, 0.94, 0.96, 0.98, and 1 are shown. (C) The proportion of all variants that are in LD with an AGV at a given threshold per ancestry group, stratified by MAF class, which is indicated by line type. The same LD thresholds from **B** are shown here.

We considered variants present in the Allen Ancient DNA Resource (Mallick et al., 2024) as AGVs. As of version 54.1, this dataset contains genotypes at 1,233,013 loci from up to 9,989 samples from 9,623 ancient individuals. After liftOver from hg19 to hg38 and excluding Y chromosome sites, which are not present in TopLD, we considered 1,199,828 total variants from the AADR (Fig. 1A).

We identified a total of 260,732,675 unique AGV-modern variant pairs at *R*^*2*^ ≥ 0.2 among all four ancestry groups in TopLD. Within ancestry groups, the number of LD variants ranged from 57,871,371 to 100,776,916. Africans had the fewest variant pairs, and Europeans had the most. This observation is consistent with expectations—Europeans have the largest sample size of all four groups and Africans exhibit more genetic diversity than non-Africans (Campbell and Tishkoff, 2008) resulting in fewer linked variants. Lower genetic diversity in non-Africans also explains the intermediate number of LD variants among East and South Asians, despite smaller sample size than Africans.

### Many common variants are in LD with variants genotyped in ancient individuals

To explore the potential of using an AGV as a proxy for a modern variant of interest, we calculated the proportion of AGV-containing variant pairs at increasing LD thresholds. We anticipated that non-African ancestry groups would have higher proportions than the African ancestry group due to lower genetic diversity in non-Africans. We stratified variant pairs by the minor allele frequency of the linked variant and strength of LD because we anticipated that the proportion of variants linked to an AGV would be positively associated with allele frequency. We classified each variant pair into one of four minor allele frequency classes: common, low-frequency, rare, or ultra-rare (Fig. 1A).

Most common variants (≥ 88%) in the four ancestry groups were linked to an AGV at the lowest threshold of *R*^*2*^ ≥ 0.2 (Fig. 1B, Table S1). As expected, non-African ancestry groups had higher proportions of common variants linked to an AGV (range = 0.952–0.958) than the African ancestry group (0.88) at a minimal *R*^*2*^. Even at a stricter threshold of *R*^*2*^ ≥ 0.9, more consistent with occurrence on the sample haplotype, ≥ 60% of modern variants in non-European ancestry grouped were linked to an AGV and 34% of African modern variants were linked. In line with expectations, variants with a MAF ≤ 5% were less likely to have high LD with an AGV (Fig. 1C). Yet, ≥ 23.3% of non-African low-frequency variants and 7.4% of African low-frequency variants were in LD with an AGV at *R*^*2*^ ≥ 0.9 (Table S1).

Rare and ultra-rare variants were the least likely to occur in LD with an AGV. Nonetheless, each ancestry group has hundreds of thousands to millions of variants in high LD with AGVs (Table S1).

### High-coverage ancient genomes confirm the co-occurrence of ancient genotyped variants and linked modern variants in recent human history

Next, we tested whether the LD between AGVs and linked variants in our catalog could be used to infer ancient genotypes at the linked variants. We examined genotypes in two high-coverage ancient human samples: Loschbour, an 8,000 year-old sample from Luxembourg (Lazaridis et al., 2014), and Ust’-Ishim, a 45,000 year-old sample from Western Siberia (Fu et al., 2014). These samples enable evaluation at different time points in human history as well as portability across lineages as the Loschbour individual contributed to modern human populations, whereas Ust’-Ishim did not (Fu et al., 2014; Lazaridis et al., 2014). We retrieved high-quality genotypes from these individuals and quantified the proportion of genotypes for AGV-modern variant pairs in the European ancestry-group that were correctly predicted at varying LD thresholds (Fig. 2A).

**Fig. 2.**
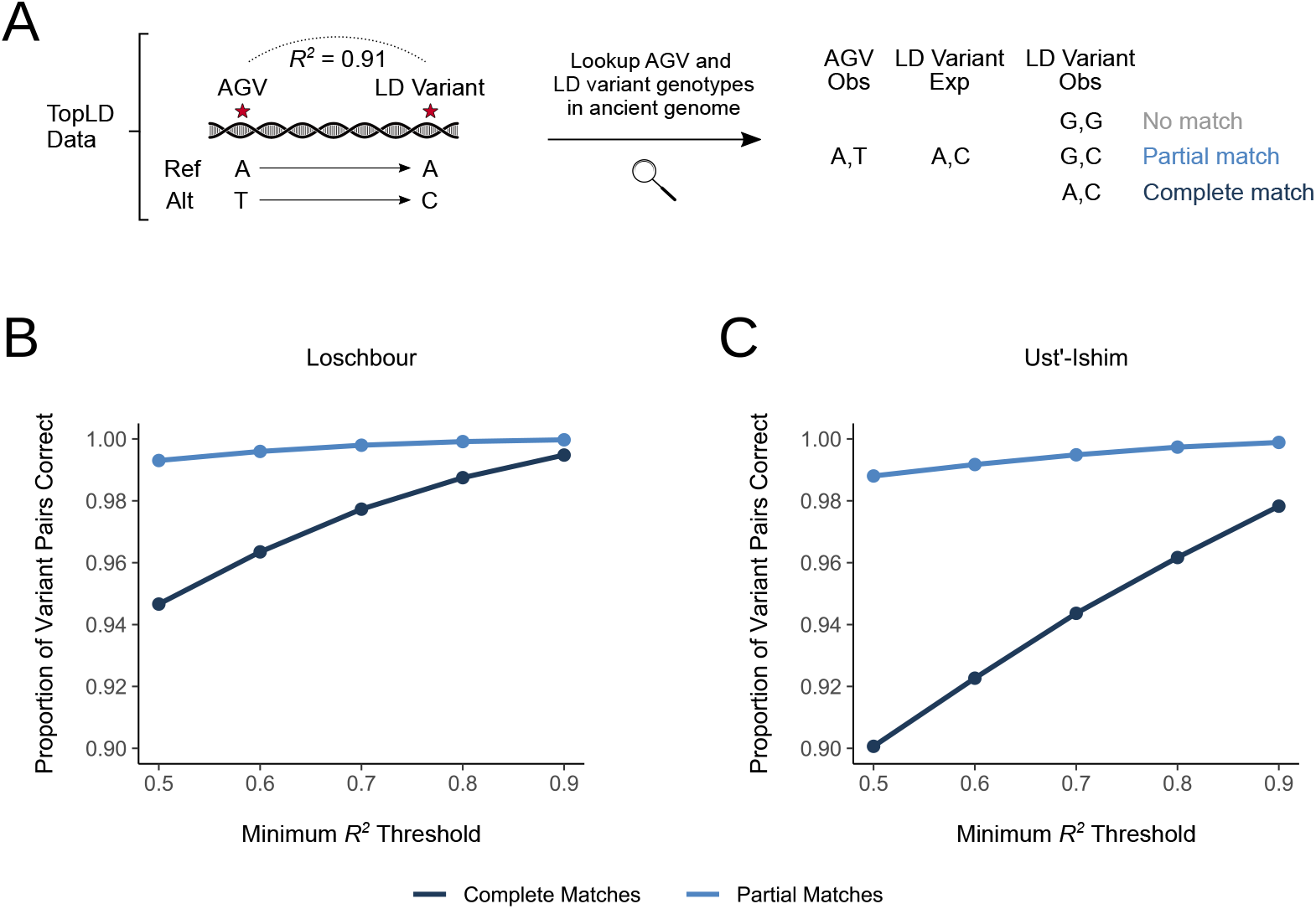
High-coverage ancient genomes confirm the co-occurrence of AGVs and modern variants in recent human history. (A) Overview of the evaluation of genotypes inferred from AGVs. We retrieved pairs of AGVs and modern variants, referred to as LD variants in this figure, in the European ancestry group with *R*^*2*^ ≥ 0.5 and mapped the expected alleles based on the correlation. Next, we retrieved high-quality genotypes from two high-coverage ancient genomes and evaluated the expected genotypes based on the associations in modern Europeans. We counted partial matches as those genotypes were one expected allele was present and complete matches as those were both expected alleles were present. We considered LD thresholds of *R*^*2*^ ≥ 0.5, 0.6, 0.7, 0.8, and 0.9. (B) Proportion of AGV-modern variant pairs partially and completely correct at varying LD thresholds for high-quality genotypes from Loschbour. Table S2 displays the number of AGV-modern variant pairs evaluated and match rates per LD threshold. Color indicates the match type. (C) Proportion of AGV-modern variant pairs partially and completely correct at varying LD thresholds for high-quality genotypes from Ust’-Ishim. Data are visualized as in **B**.

Greater than 99% of predicted co-occurring AGVs and modern variants at high LD (*R*^*2*^ ≥ 0.9) were partially or completely correct in genotypes from Loschbour (Fig. 2B). Partial matches also occurred at a similar proportion at a modest LD threshold (*R*^*2*^ ≥ 0.5) and the complete match rate was 94.7%. In an older and more distantly genetically related sample—Ust’-Ishim, European AGV-modern variant pairs were also robustly predicted. 97.8% of pairs at high LD occurred as complete matches in Ust’-Ishim (Fig. 2C). Even at a modest LD threshold, 90% of pairs were correctly called as complete matches in this older sample.

These results support the use of the AGV-modern variant pairs curated here to rapidly identify variants with ancient genotyping data that co-occur with individual variants of interest without the need for complex imputation.

### Most ancient genotyped variants are present among all TopLD ancestry groups

Next, we characterized the presence of AGVs in TopLD. 1,162,596 or 94.3% of AGVs were present in TopLD, while 69,890 were absent. We predicted that absent variants would be largely explained by low global allele frequencies. Indeed, AGVs absent from TopLD have a median allele frequency of 3.94 *×* 10^5^, whereas those present have a median allele frequency of 0.237 (Mann-Whitney, U = 1,983,228,367.5, P = 2.23 *×* 10^308^) (Fig. S1). However, we still observed 3,090 variants with an allele frequency ≥ 0.01 that were absent, including several high frequency variants.

We further examined the distribution of AGVs among the four ancestry groups. As expected, most AGVs (N = 944,223) were shared among all four groups (Fig. S2). The next most abundant were AGVs shared by Africans and Europeans (N = 110,863). This pattern is likely explained by the higher power in those groups due to higher sample sizes. Among unique AGVs, Africans had the highest (N = 25,725), followed by Europeans (N = 4,795), East Asian (N = 3,175), and South Asians (N = 603). Allele frequency distributions are largely similar across ancestry groups; however, Africans have an increased proportion of higher frequency variants (Fig. S3). These patterns are expected due to human evolutionary history and sampling in TopLD.

### Ancient genotyped variants are significantly older than expected

We quantified the distribution of AGV ages by retrieving age estimates from the Human Genome Dating project (Albers and McVean, 2020). Estimated ages ranged from 625 to 2,192,115 years, with a median age of 695,106 years—assuming a generation time of 29 years (Fenner, 2005). We compared this distribution to a distribution comprised of age estimates for random variants, matching on allele frequency bins and estimation method (see Methods). Randomly-sampled variants were significantly younger than AGVs with median age of 644,493 years; > 50,000 years younger than the AGVs (Mann-Whitney, U = 470,998,932,171, P = 2.23 *×* 10^308^) (Fig. 3).

**Fig. 3.**
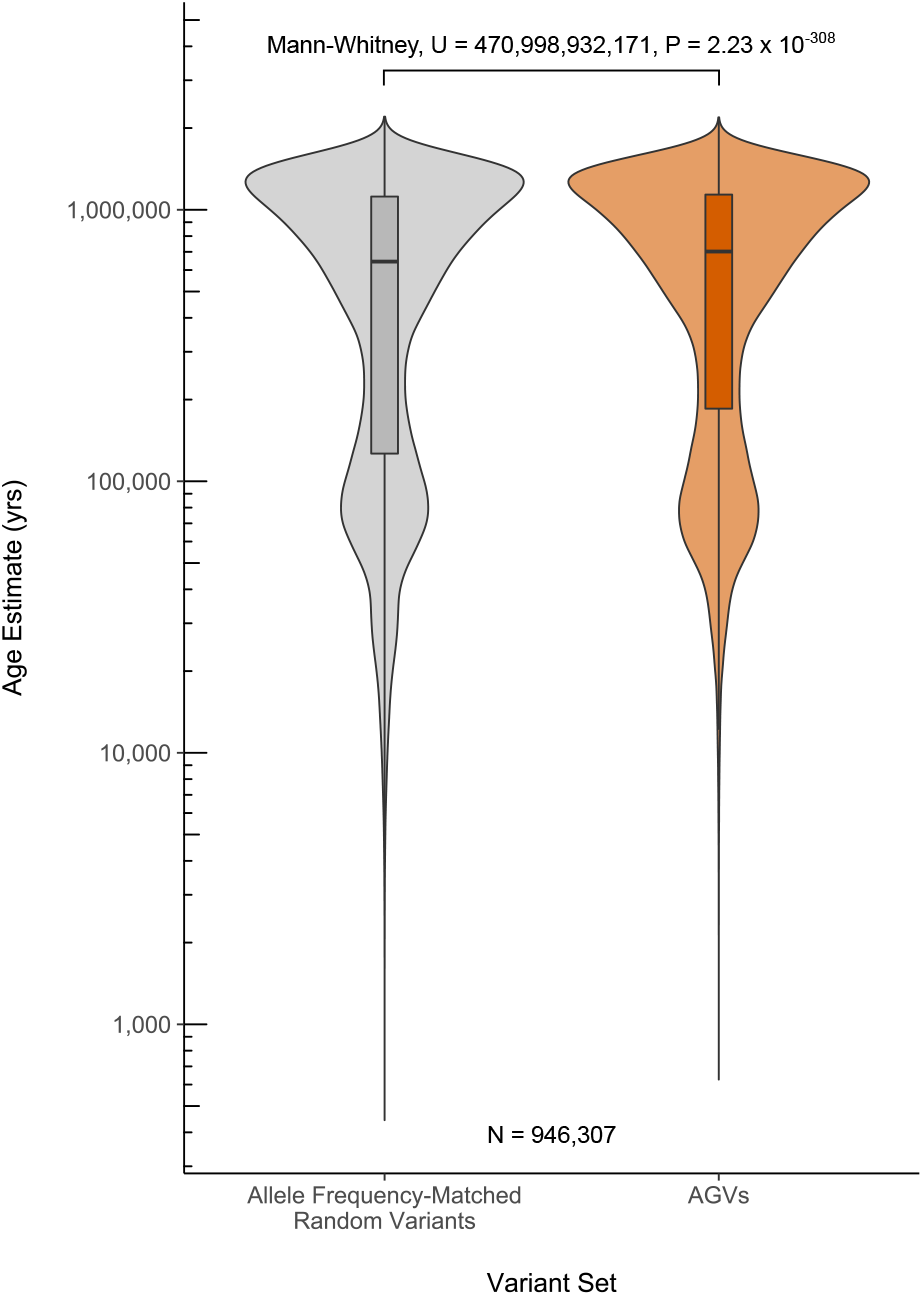
Ancient genotyped variants are significantly older than expected. The estimated allele age distribution for 946,307 high-quality AGVs compared to source- and allele-frequency matched random variants. We used the mode age estimated using the joint clock per variant from the Human Genome Dating project (Albers and McVean, 2020).

This is likely due to the design of the capture arrays used that focus on variants with low to common minor allele frequencies across multiple human populations. Such patterns are expected from variants that evolved before or shortly after humans migrated out of Africa, ∼ 60,000 years ago.

### Ancient-genotyped variants are characterized by stable allele frequencies over time

Haplotypes that have recently arisen or experienced recent positive selection are of both clinical and evolutionary interest. Only 77,686 or 8.1% of AGVs are estimated to be ≤ 60,000 years old and to have presumably appeared after the ancestors of modern humans migrated out of Africa. Further, the oldest modern human sample present in the AADR is 45,371 years old. Thus, the ability to capture the origin of individual alleles is restricted to a subset of ancient genotyped variants.

Among these variants, only a fraction are expected to have experienced recent non-neutral evolution. However, the dynamics of older haplotypes may also be detected if such selection has happened in the past 50,000 years. Therefore, despite the limited number of variants for which observing allele origin is possible, the number of variants where allele frequencies may be dynamic is potentially greater.

We illustrate the potential for our database to enable study of origin and dynamics using two examples. First, we highlight a well-characterized locus that experienced recent positive selection in some humans populations. A missense variant (rs3827760) in *EDAR* that occurs at high-frequency among individuals with genetic similarity to Asian samples from the HapMap2 Project (Sabeti et al., 2007). This allele is associated with multiple phenotypic effects in mice and humans, including increased hair thickness and an increased number of eccrine glands (Kamberov et al., 2013). The estimated age for this allele is 36,410 years and allele frequencies for this variant remain zero until exhibiting noticeable increases in ancient samples from Asia and the Americas ∼ 20,000 years ago (Fig. 4A).

**Fig. 4.**
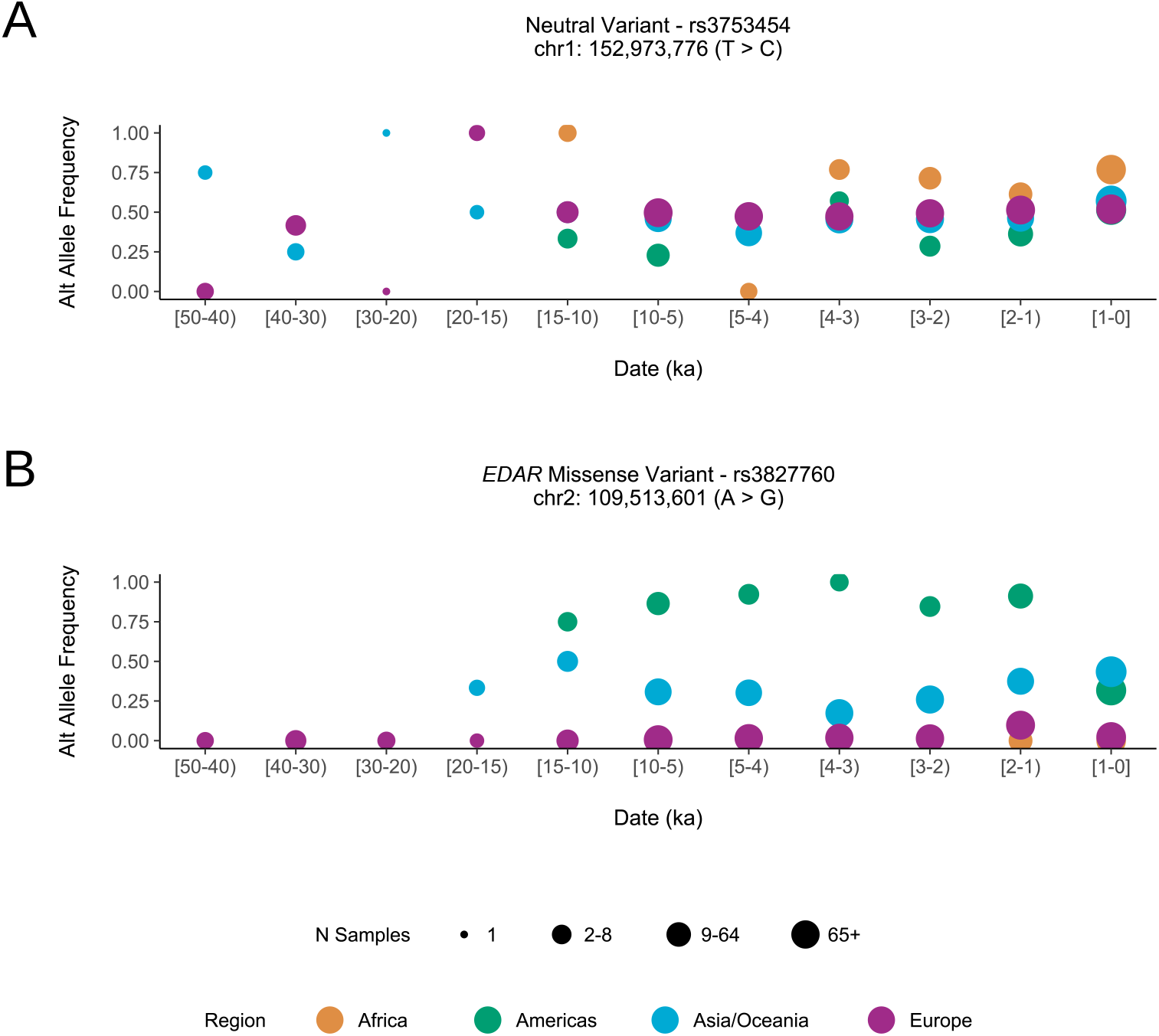
Ancient-genotyped variant allele frequencies over time. (A) Allele frequencies over time in ancient and modern humans for an *EDAR* missense variant: rs3827760 or chr2:109,513,601 (A > G). This variant is estimated to have emerged 36,410 years ago and been subject to positive selection shortly after emergence. This locus is a well-characterized target of recent selection in East Asians. Point size indicates the number of samples used to calculate allele frequency and point color indicates the geographic region. (B) Allele frequencies over time in ancient and modern humans for a putatively neutral variant: rs3753454 or chr1:152,973,776 (T > C). This variant is estimated to have emerged 933,855 years ago. Point size indicates the number of samples used to calculate allele frequency and point color indicates the geographic region. The allele frequency patterns here are typical for most AGVs and reflect a locus subject to neutral evolution across human populations. Data are visualized as in **A**.

As a second example, we analyze an example intergenic variant on chromosome 1, rs3753454, with no functional annotations, that is unlikely to have experienced strong selection. and which is estimated to have appeared 933,855 years ago (Fig. 4B). Ancient allele frequencies in Asia and Europe, the best sampled geographical regions, are stable through time, especially in the recent past where sample sizes are larger. Such patterns are consistent with a locus under neutral evolution.

### A catalog for the rapid identification of AGVs linked to modern variants

Identification of proxy AGVs for variants of interest can facilitate study of the dynamics of common variants, including those implicated in genetic association studies. To illustrate this, we used our catalog to analyze a variant, rs72823641, that is consistently associated with decreased risk of asthma (Ferreira et al., 2019; Sakaue et al., 2021) and was recently identified as a putative positively selected locus in West Eurasians (Akbari et al., 2024).

This variant is not an AGV, so it cannot be studied in ancient samples directly. However, rs72823641 is in high LD (*R*^*2*^ ≥ 0.9) with five AGVs, all of which are shared between Europeans and South Asians and in high LD with one another (Fig. S4). Using these AGVs, we can trace the frequency of the query variant over time, and observe an increase in allele frequency through the present in Europe and South Asia.

rs72823641 is also in modest LD with AGVs in other populations; however, given that the LD is comparatively low (*R*^*2*^ 0.383 in Africans, *R*^*2*^ of 0.769 in East Asians), we caution against using them as proxies. We also note that the AGVs are estimated to be considerably older than rs72823641, 122,655–151,000 years compared to 71,420 years (Albers and McVean, 2020). Thus, these variants likely emerged after modern humans diverged from the shared lineage with Neanderthals—the alternate alleles are absent in all three high-coverage Neanderthals. This suggests that rs72823641 may have emerged on this haplotype following humans’ last major emigration from Africa.

To enable similar analyses, we provide a tool that researchers can use to identify whether or not a variant of interest is an AGV or has an AGV in strong LD that could serve as a proxy (https://github.com/brandcm/ancient_genotyped_variants_proxy_catalog_app). Users can specify genomic coordinates in hg38 or use an rsID. If a variant is an AGV, users are provided with allele frequencies from ancient samples, as well as genotypes from four high-coverage archaic hominins. Both proxy variants and genotypes for AGVs are available for download for a locus of interest or as bulk data.

## DISCUSSION

Here, we identify modern variants in LD with commonly genotyped variants in ancient individuals. We then evaluate how often such variants can serve as proxies for variants of interest. These analyses expand the utility of existing ancient genomes in the study of allele histories and enable analysis of many variants beyond those directly genotyped in ancient samples.

We find that the majority of common variants in non-Africans are in strong LD with at least one AGV. In Africans, proportionally fewer variants are in LD with AGVs, but millions of African variants are in strong LD with at least one AGV. Further, many variants at low allele frequencies also occur with AGVs, potentially enabling study of their origins. We tested the ability to infer genotypes through time based on LD with AGVs using two high-coverage samples. The proxy variants yielded accurate predicted genotypes (> 90% complete match at LD above 0.5), even in a 45,000 year old sample that did not directly contribute to modern European populations. Collectively, our results reveal that many modern variants of interest can be studied in the recent past by using AGV proxies.

We also characterized the AGVs themselves, revealing that many are present among different ancestry groups, and that they are older than expected. This, combined with the small number of ancient human samples from > 20,000 years ago (N = 17), limits the ability to observe allele birth from AGVs. However, it is possible to trace dynamics in recent allele frequencies, which can be informative when studying the presence and timing of recent selection.

To enable more researchers to incorporate ancient DNA into their analyses, we provide a website for querying whether modern variants of interest are AGVs and a database of modern variants linked to AGVs. This resource enables rapid identification of any linked variants for variants of interest that are not AGVs themselves without the need for imputation. More complex genotype imputation methods will likely be useful in ancient DNA applications, but they are computationally intensive and dependent on appropriate reference panels (Escobar-Rodríguez and Veeramah, 2024; Garrido Marques et al., 2024; Sousa da Mota et al., 2023). Our tool aims to streamline and facilitate evolutionary analyses of variants of interest. Combining it with other recent ancient DNA tools, such as DORA (Harris and Greenbaum, 2024), which allows users to visualize ancient samples and associated metadata geographically, we anticipate that a larger community will be able to make use of this powerful resource.

We acknowledge several limitations of our tool and approach. While millions of modern variants are strongly linked to AGVs, many variants of interest, especially rare variants, may not be linked to an AGV. In addition, new ancient DNA data are released regularly, and while the AADR is by far the largest, our tool does not integrate all ancient samples. For modern variants, we used the ancestry groups present in the TopLD dataset for our analyses; however, these panels may not be suitable for understanding all variants of interest in more specific populations. Finally, the history of the linkage landscape for haplotypes is quite complex, as new mutations are added. Thus, age estimation of variants of interest themselves using resources such as the Human Genome Dating project are informative for understanding haplotype history (Yee et al., 2024).

As increasing numbers of variants are implicated in human health and disease from association tests and other methods, our tool enables researchers to quickly gain evolutionary perspectives on variants of interest.

## METHODS

### Ancient Human Genotypes

We analyzed AGVs from the Allen Ancient DNA Resource, version 54.1.p1 (Mallick et al., 2024). We used liftOver (Hinrichs et al., 2006) to translate AGV hg19 coordinates to the hg38 reference genome. This process removed 527 variants absent in the hg38 assembly. We also switched the reference and alternate alleles for 3,676 variants using a custom script due to reference sequence differences between assemblies. Finally, we did not consider the 32,658 Y chromosome AGVs or 140 AGVs that mapped to alternate scaffolds on chromosomes 7, 8, and 19 due to lack of LD data. This resulted in 1,199,828 total AGVs considered in downstream linked variant analyses.

### Modern Human Variants in Linkage Disequilibrium with AGVs

We used pairwise variant LD information from TopLD (Huang et al., 2022; Taliun et al., 2021). These data were derived from autosomal and X chromosome WGS data in four TOPMed cohorts: BioMe, MESA, JHS, and WHI. Local and global ancestry were estimated per sample using reference populations from the Thousand Genomes Project (TGP; Auton et al., 2015) and the Human Genome Diversity Project (HGDP; Rosenberg et al., 2002). Global ancestry was used to categorize unrelated individuals into one of four “ancestry groups”: African, East Asian, European, and South Asian. LD was then estimated separately in each population for all variant pairs within 500 kb, and variant pairs with *R*^*2*^ ≥ 0.2 were retained. We retrieved variant pairs from each ancestry group dataset that included an AGV. We used the minor allele frequency (MAF) provided in the TopLD annotation files to quantify the frequencies of TopLD variants in LD with an AGV. Variants were classified as common (MAF = >0.05), low-frequency (MAF = [0.01–0.05)), rare (MAF = [0.001–0.01)), and ultra-rare (< 0.001).

### Evaluation in High-Coverage Ancient Genomes

We retrieved high-quality variants from two high-coverage ancient genomes: Loschbour, an 8,000 year old sample from the Loschbour rock shelter in Luxembourg (Lazaridis et al., 2014), and Ust’-Ishim, a 45,000 year old sample from southern Siberia (Fu et al., 2014). We restricted variants to those that passed a suite of filters used to process several high-coverage ancient samples by the Max Planck Institute for Evolutionary Anthropology, including positions that do not overlap indels, fall within a tandem repeat, or whose mapping quality is < 25 (Prüfer et al., 2017). We further set genotypes whose coverage was < 10 and whose genotype quality < 30 to missing and removed these genotypes using bcftools, version 1.21 (Li, 2011). Next, we used liftOver to convert positions from hg19 to hg38 (Hinrichs et al., 2006). This resulted in 1,652,392,042 genotyped loci for Loschbour and 1,769,322,914 genotyped loci for Ust’-Ishim.

To evaluate the proportion of correctly predicted genotypes in each ancient sample, we focused on 61,485,131 AGV-modern variant pairs from the European ancestry group with *R*^*2*^ ≥ 0.5. We generated a predicted allele set at the linked modern variant site for each AGV-modern variant pair. Given that indels are not called in these ancient samples, we excluded any AGV-modern variant pairs where either the reference or alternate allele were longer than one nucleotide. We required that both genotypes for both the AGV and modern variant were present in the evaluated ancient genome. For positively correlated pairs, the AGV reference allele co-occurred with the modern variant reference allele as well as the AGV alternate allele co-occurring with the modern variant alternate allele. For negatively correlated pairs, the AGV reference allele co-occurred with the modern variant alternate allele and vice-versa. Based on the count of AGV reference and alternate alleles, we generated the predicted allele set. Each AGV-modern variant pair was classified as a “no match” if none of the predicted alleles were present in the evaluation genome, a “partial match” if only one was present, and a “complete match” if both were present. Table S2 displays the number of AGV-modern variant pairs evaluated and match rates per LD threshold per sample.

### Ancient Genotyped Variant Age Estimates

We retrieved allele age estimates for AGVs from the Human Genome Dating project (Albers and McVean, 2020). This dataset comprises age estimates for autosomal variants identified in samples from the TGP (Auton et al., 2015) and the Simons Genome Diversity Project (SGDP) (Mallick et al., 2016). Briefly, allele ages are estimated using a non-parametric approach, where an ancestral genomic segment relative to a target variant is identified using a hidden Markov model. The number of mutations and genetic length of the segment is then measured between pairs of sample chromosomes, which are fed into probabilistic models that estimate allele age. Three “clocks” were used to estimate age: 1) a clock that uses the number of mutations or “mutation clock”, 2) a clock that uses ancestral segment’s length or “recombination clock”, and 3) a clock that uses both the number of mutations and genetic length or “joint clock”. For each clock, the mean, median, mode, and lower & upper bounds of the 95% confidence interval of the composite posterior probability are reported. Variants private to either SGDP or TGP are estimated using solely those data. However, variants present in both data sources were analyzed using each dataset separately as well as a combined dataset. Therefore, each variant had up to three age estimates from one of three data sources: combined, SGDP, and TGP.

We intersected our set of AGVs with variants in the Human Genome Dating project, retrieving 3,126,750 age estimates for 1,117,110 AGVs. We verified that quality scores were not associated with allele age in any of the estimation methods among AGVs (Fig. S5). Quality score distribution was also comparable across estimation methods, although estimates using the recombination clock were more likely to have lower quality scores (Fig. S5). We choose to use the age estimates from the joint clock in this analysis to maximize the information derived from sample haplotypes. Following the recommendation of the authors, we used the mode reported per variant from the joint clock as the per variant age estimate.

2,294,708 high quality age estimates for 954,794 AGVs remained after excluding low quality estimates per author recommendations (quality score ≤ 0.5). We also compared the age estimates across high-quality variants with age estimates from all three data sources (N = 637,502) to test how data source affected age estimates. We note that other factors, largely the number of haplotypes with a given variant, also impacts age estimation under this method. Age estimates derived from SGDP data were slightly younger (*μ* = 638,776 years) compared to those derived from the combined dataset (*μ* = 691,410 years) and TGP (*μ* = 697,328 years) (Fig. S6). These distributions are significantly different (Kruskal-Wallis, H = 4066.34, P = 2.23 *×* 10^308^); however, the effect size between estimates derived from combined data compared to those derived from TGP data is small (Fig. S6).

We generated a distribution of AGV allele ages using a single estimate per variants, prioritizing estimates from combined data sources where possible and mapped global allele frequencies to each variant using data from gnomAD, version 2.1.1 (Karczewski et al., 2020) for variants whose “FILTER” was equal to “PASS”. This resulted in age estimates for 946,307 AGVs where combined data comprised 697,199 estimates, SGDP data comprised 179,130 estimates, and TGP data comprised 69,978 estimates. We compared this distribution to a random sample (N = 946,307) of variant ages from the Human Genome Dating project. We matched each AGV with a random variant on 100 allele frequency bins (e.g., (0.02-0.03]) and maintained the distribution of estimates from combined, SGDP, and TGP data sources. We then compared the distribution of ages for AGVs and the random variants using a Mann-Whitney U-test.

### Allele Frequency Trajectories

We retrieved allele frequencies for variants of interest from the AADR. Each sample was categorized into a geographic region based on the country in which the sample was found: Africa, Americas, Asia, Europe, and Oceania. We excluded genotypes from any reference sequence or archaic hominins as well as missing genotypes. For samples with multiple genotype calls, we prioritized the “Shotgun diploid” data source when available to capture heterozygous genotypes. If this data was not present for a sample with multiple data sources, we used the first data source. We binned samples into one of 11 non-overlapping time periods in ka: [50-40), [40-30), [30-20), [20-15), [15-10), [10-5), [5-4), [4-3), [3-2), [2-1), and [1-present).

### Analysis

All data analyses were performed using Bash and Python scripts, some of which were implemented in Jupyter notebooks. All reported p-values are two-tailed, unless noted otherwise.

### Visualization

Results were visualized using Inkscape, version 1.1 (Inkscape Project, 2020) and ggplot, version 3.3.6 (Wickham, 2016) implemented in R, version 4.0.5 (R Core Team, 2020).

## Data and Code Availability

We used publicly available data for all analyses. Ancient genotyped variants were retrieved from https://reich.hms.harvard.edu/allen-ancient-dna-resource-aadr-downloadable-genotypes-present-day-and-ancient-dna-data. Modern variant LD data were retrieved from TopLD (http://topld.genetics.unc.edu/downloads/downloads/). Genotypes from high-coverage ancient human genomes were retrieved from http://cdna.eva.mpg.de/neandertal/Vindija/VCF/. Allele age estimates were retrieved from the Human Genome Dating project (https://human.genome.dating/download/index).

All code used to conduct analyses and generate figures is publicly available on GitHub (https://github.com/brandcm/ancient_genotyped_variants_proxy_catalog).

## Acknowledgements

We thank members of the Capra Lab who gave helpful feedback throughout this project. We are grateful for the support and resources from the Wynton High Performance Compute Cluster at University of California San Francisco. JAC and CMB were funded by National Institutes of Health grant R35GM127087.

## Author Contributions

Conceptualization, CMB and JAC; Formal Analysis, CMB; Writing – Original Draft, CMB and JAC; Writing – Review & Editing, CMB and JAC.

## References

Akbari A. et al. 2024. Pervasive Findings of Directional Selection Realize the Promise of Ancient DNA to Elucidate Human Adaptation. bioRxiv: 2024.09.14.613021. DOI: 10.1101/2024.09.14.613021.

Albers P. K. and McVean G. 2020. Dating Genomic Variants and Shared Ancestry in Population-Scale Sequencing Data. PLOS Biology 18:e3000586. DOI: 10.1371/journal.pbio.3000586.

Auton A. et al. 2015. A Global Reference for Human Genetic Variation. Nature 526: 68–74. DOI: 10.1038/nature15393.

Barrie W., Irving-Pease E. K., Willerslev E., Iversen A. K. N., and Fugger L. 2024. Ancient DNA Reveals Evolutionary Origins of Autoimmune Diseases. Nature Reviews Immunology 24: 85–86. DOI: 10.1038/s41577-023-00983-6.

Campbell M. C. and Tishkoff S. A. 2008. African Genetic Diversity: Implications for Human Demographic History, Modern Human Origins, and Complex Disease Mapping. Annual Review of Genomics and Human Genetics 9: 403–433. DOI: 10.1146/annurev.genom.9.081307.164258.

Escobar-Rodríguez M. and Veeramah K. R. 2024. Evaluation of Ancient DNA Imputation: A Simulation Study. Human Population Genetics and Genomics 4: 0002. DOI: 10.47248/hpgg2404010002.

Fenner J. N. 2005. Cross-Cultural Estimation of the Human Generation Interval for Use in Genetics-Based Population Divergence Studies. American Journal of Physical Anthropology 128: 415–423. DOI: 10.1002/ajpa.20188.

Ferreira M. A. et al. 2019. Genetic Architectures of Childhood- and Adult-Onset Asthma Are Partly Distinct. The American Journal of Human Genetics 104: 665–684. DOI: 10.1016/j.ajhg.2019.02.022.

Fu Q. et al. 2014. Genome Sequence of a 45,000-Year-Old Modern Human from Western Siberia. Nature 514: 445–449. DOI: 10.1038/nature13810.

Garrido Marques A., Rubinacci S., Malaspinas A.-S., Delaneau O., and Sousa da Mota B. 2024. Assessing the Impact of Post-Mortem Damage and Contamination on Imputation Performance in Ancient DNA. Scientific Reports 14: 6227. DOI: 10.1038/s41598-024-56584-3.

Harris K. D. and Greenbaum G. 2024. DORA: An Interactive Map for the Visualization and Analysis of Ancient Human DNA and Associated Data. Nucleic Acids Research 52:W54– W60. DOI: 10.1093/nar/gkae373.

Hinrichs A. S. et al. 2006. The UCSC Genome Browser Database: Update 2006. Nucleic Acids Research 34: D590–D598. DOI: 10.1093/nar/gkj144.

Huang L. et al. 2022. TOP-LD: A Tool to Explore Linkage Disequilibrium with TOPMed Whole-Genome Sequence Data. The American Journal of Human Genetics 109: 1175–1181. DOI: 10.1016/j.ajhg.2022.04.006.

Inkscape Project. 2020. Inkscape.

Kamberov Y. G. et al. 2013. Modeling Recent Human Evolution in Mice by Expression of a Selected EDAR Variant. Cell 152: 691–702. DOI: 10.1016/j.cell.2013.01.016.

Karczewski K. J. et al. 2020. The Mutational Constraint Spectrum Quantified from Variation in 141,456 Humans. Nature 581: 434–443. DOI: 10.1038/s41586-020-2308-7.

Lazaridis I. et al. 2014. Ancient Human Genomes Suggest Three Ancestral Populations for Present-Day Europeans. Nature 513: 409–413. DOI: 10.1038/nature13673.

Li H. 2011. A Statistical Framework for SNP Calling, Mutation Discovery, Association Mapping and Population Genetical Parameter Estimation from Sequencing Data. Bioinformatics 27: 2987–2993. DOI: 10.1093/bioinformatics/btr509.

Mallick S., Micco A., Mah M., Ringbauer H., Lazaridis I., Olalde I., Patterson N., and Reich D. 2024. The Allen Ancient DNA Resource (AADR) a Curated Compendium of Ancient Human Genomes. Scientific Data 11: 182. DOI: 10.1038/s41597-024-03031-7.

Mallick S. et al. 2016. The Simons Genome Diversity Project: 300 Genomes from 142 Diverse Populations. Nature 538: 201–206. DOI: 10.1038/nature18964.

Mathieson S. and Mathieson I. 2018. FADS1 and the Timing of Human Adaptation to Agriculture. Molecular Biology and Evolution 35: 2957–2970. DOI: 10.1093/molbev/msy180.

Orlando L. et al. 2021. Ancient DNA Analysis. Nature Reviews Methods Primers 1: 14. DOI: 10.1038/s43586-020-00011-0.

Pääbo S., Poinar H., Serre D., Jaenicke-Després V., Hebler J., Rohland N., Kuch M., Krause J., Vigilant L., and Hofreiter M. 2004. Genetic Analyses from Ancient DNA. Annual Review of Genetics 38: 645–679. DOI: 10.1146/annurev.genet.37.110801.143214.

Patin E. and Quintana-Murci L. 2024. Tracing the Evolution of Human Immunity through Ancient DNA. Annual Review of Immunology. DOI: 10.1146/annurev-immunol-082323-024638.

Prüfer K. et al. 2017. A High-Coverage Neandertal Genome from Vindija Cave in Croatia. Science 358: 655–658. DOI: 10.1126/science.aao1887.

R Core Team. 2020. R: A Language and Environment for Statistical Computing. R Foundation for Statistical Computing. Vienna, Austria.

Rosenberg N. A., Pritchard J. K., Weber J. L., Cann H. M., Kidd K. K., Zhivotovsky L. A., and Feldman M. W. 2002. Genetic Structure of Human Populations. Science 298: 2381–2385. DOI: 10.1126/science.1078311.

Sabeti P. C. et al. 2007. Genome-Wide Detection and Characterization of Positive Selection in Human Populations. Nature 449: 913–918. DOI: 10.1038/nature06250.

Sakaue S. et al. 2021. A Cross-Population Atlas of Genetic Associations for 220 Human Phe-notypes. Nature Genetics 53: 1415–1424. DOI: 10.1038/s41588-021-00931-x.

Slatkin M. and Racimo F. 2016. Ancient DNA and Human History. Proceedings of the National Academy of Sciences 113: 6380. DOI: 10.1073/pnas.1524306113.

Sousa da Mota B. et al. 2023. Imputation of Ancient Human Genomes. Nature Communications 14: 3660. DOI: 10.1038/s41467-023-39202-0.

Taliun D. et al. 2021. Sequencing of 53,831 Diverse Genomes from the NHLBI TOPMed Program. Nature 590: 290–299. DOI: 10.1038/s41586-021-03205-y.

Wickham H. 2016. Ggplot2: Elegant Graphics for Data Analysis. New York: Springer-Verlag. ISBN: 978-3-319-24277-4.

Willerslev E. and Cooper A. 2004. Review Paper. Ancient DNA. Proceedings of the Royal Society B: Biological Sciences 272: 3–16. DOI: 10.1098/rspb.2004.2813.

Yee S. W. et al. 2024. Illuminating the Function of the Orphan Transporter, SLC22A10, in Humans and Other Primates. Nature Communications 15: 4380. DOI: 10.1038/s41467-024-48569-7.

Yilmaz F. et al. 2025. Reconstruction of the Human Amylase Locus Reveals Ancient Duplications Seeding Modern-Day Variation. Science 386 ():eadn0609. DOI: 10.1126/science.adn0609.

