## Supplemental Information for "A catalog of ancient proxies for modern genetic variants"

### Supplementary Information

#### Supplemental Figures

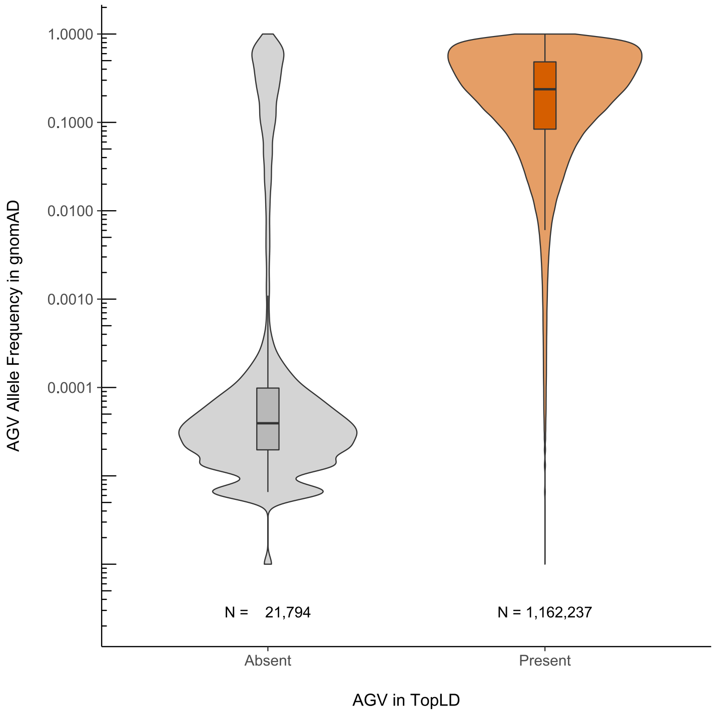


**Fig. S1. AGVs absent from TopLD are largely rare and ultra-rare variants.** The distribution of global allele frequencies from gnomAD for AGVs, stratified by absence/presence in TopLD. Note the y-axis is log10 transformed. Distributions are visualized as boxplots where the boxes indicate the median and IQR, with the upper whiskers extending to the largest value $\leq$ 1.5 x IQR from the 75th percentile and the lower whisksers extending to the smallest value $\leq$ 1.5 x IQR from the 25th percentile.

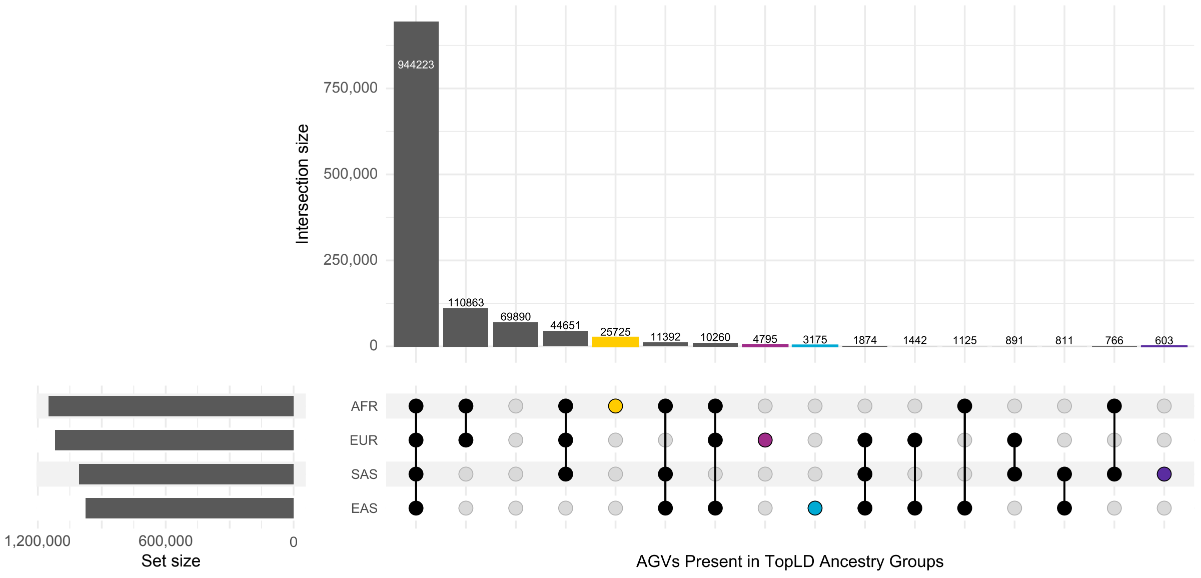


**Fig. S2. Unique and shared AGVs in TopLD among ancestry groups.** Unique and shared AGVs present in TopLD and in LD $R^2 \geq 0.2$ with at least one modern variant among ancestry groups. The bars indicate the size of each group and the dot matrix indicates the populations comprising each set.

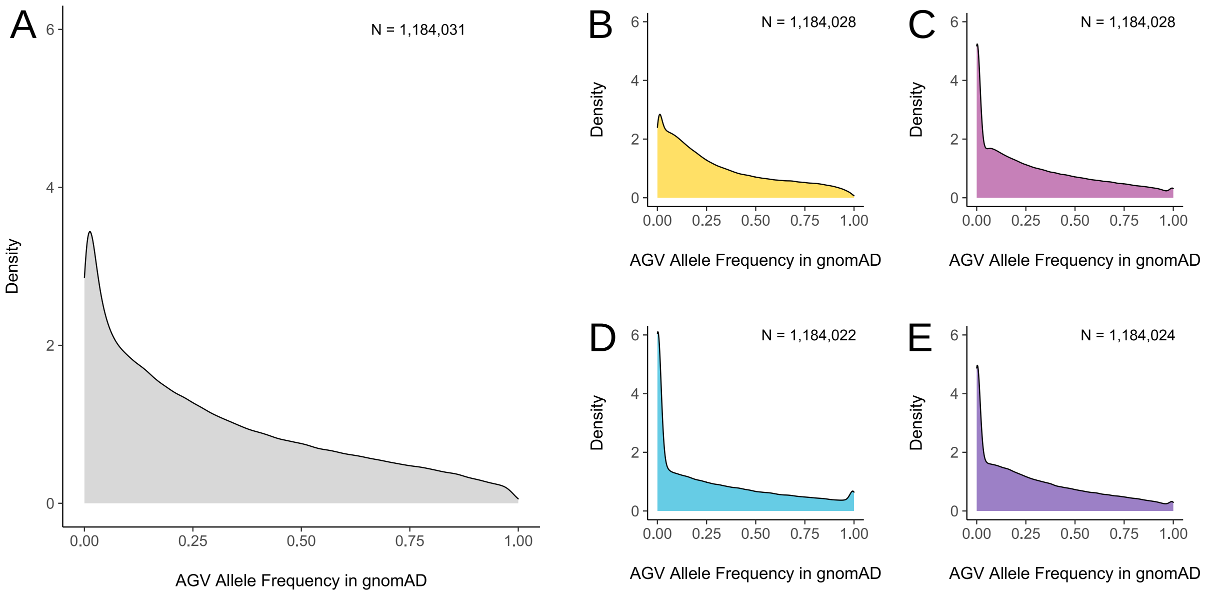


**Fig. S3. AGVs allele frequencies by ancestry group.** The total allele frequency distribution in gnomAD for AGVs (A) and stratified by ancestry groups: African/African-American (B), East Asian (C), European (D), and South Asian (D).

AGVs in LD with Variant of Interest

| Variant of Interest | | | | | Ancient Genotyped Variant | | | | LD Information | | | |
|---|---|---|---|---|---|---|---|---|---|---|---|---|
| Chromosome | Position | Ref Allele | Alt Allele | rsID | Position | Ref Allele | Alt Allele | rsID | Ancestry Groups | r² | D' | Correlation |
| 2 | 102319699 | T | A | rs72823641 | 102337157 | G | T | rs3771180 | EUR,SAS | 0.974,1.0 | 0.994,1.0 | +,+ |
| 2 | 102319699 | T | A | rs72823641 | 102338622 | G | A | rs13408661 | EUR,SAS | 0.974,1.0 | 0.994,1.0 | +,+ |
| 2 | 102319699 | T | A | rs72823641 | 102338193 | C | T | rs13431828 | EUR,SAS | 0.971,1.0 | 0.991,1.0 | +,+ |
| 2 | 102319699 | T | A | rs72823641 | 102343750 | T | A | rs3771175 | EUR,SAS | 0.965,1.0 | 0.991,1.0 | +,+ |
| 2 | 102319699 | T | A | rs72823641 | 102350089 | A | G | rs10197862 | EUR,SAS | 0.961,1.0 | 0.991,1.0 | +,+ |
| 2 | 102319699 | T | A | rs72823641 | 102270668 | T | C | rs11674302 | EUR,SAS | 0.875,0.757 | 0.949,0.899 | +,+ |
| 2 | 102319699 | T | A | rs72823641 | 102267515 | C | T | rs11692065 | EUR,SAS | 0.874,0.757 | 0.948,0.899 | +,+ |
| 2 | 102319699 | T | A | rs72823641 | 102263004 | C | A | rs10189629 | EUR,SAS | 0.873,0.757 | 0.948,0.899 | +,+ |
| 2 | 102319699 | T | A | rs72823641 | 102250184 | G | A | rs13406331 | EUR,SAS | 0.839,0.607 | 0.92,0.793 | +,+ |
| 2 | 102319699 | T | A | rs72823641 | 102183291 | T | C | rs11895033 | EAS | 0.769 | 0.899 | + |

1 / 5

**Fig. S4. Top AGVs identified for rs72823641.** The top ten AGVs identified for rs72823641.

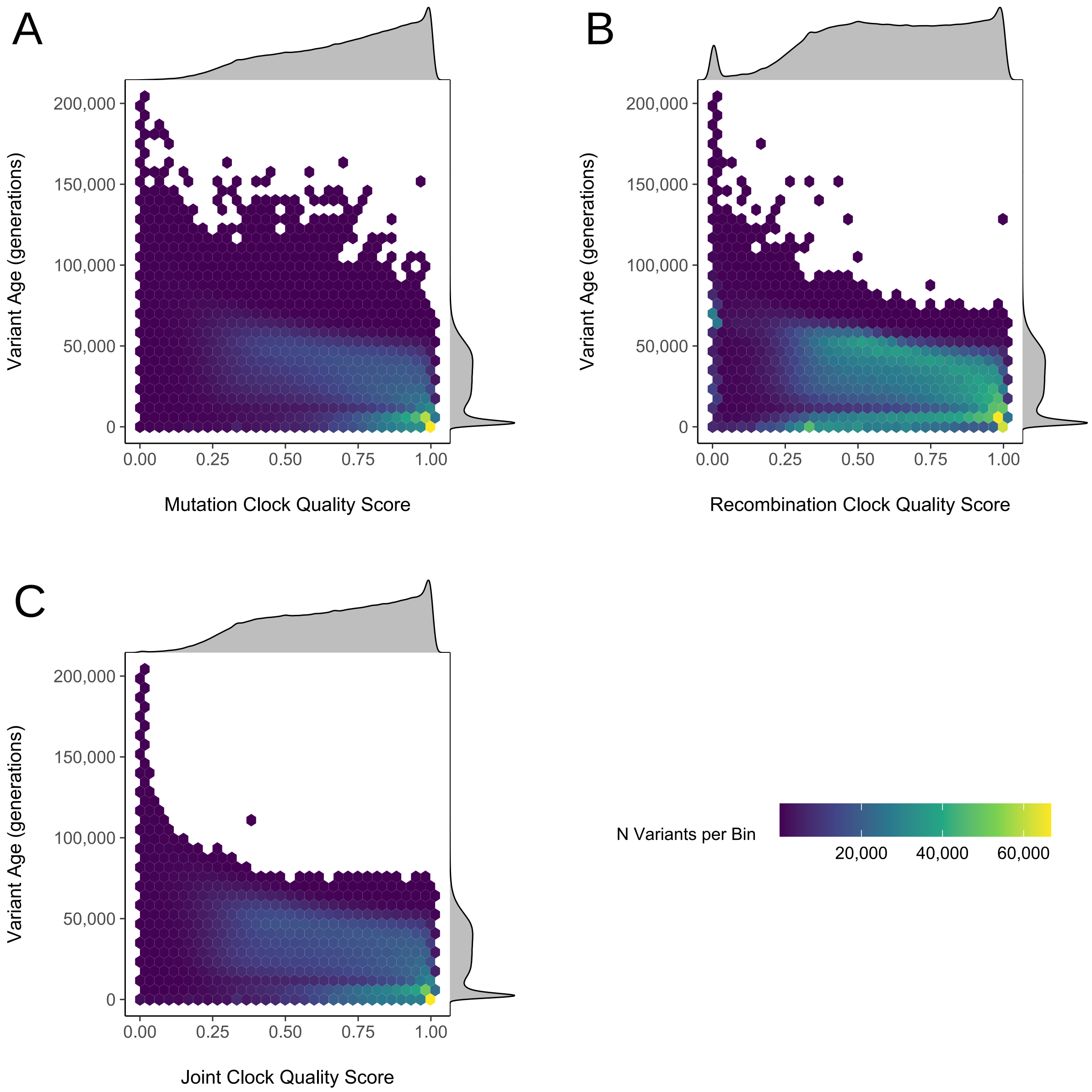


**Fig. S5. Quality scores of AGVs dated in the Human Genome Dating project are not associated with allele age.** Hex bins of clock quality score and estimated variant age in generations for AGVs dated in the Human Genome Dating project (Albers and McVean, 2020). Quality scores range from 0 indicating poor quality of the estimate to 1 indicating high quality of the estimate. The number of variants per bin in indicated by color. The individual distributions for both estimated variant age and clock quality score are shown using density plots. The mutation clock is shown in **A**, the recombination clock is shown in **B**, and the joint clock is shown in **C**.

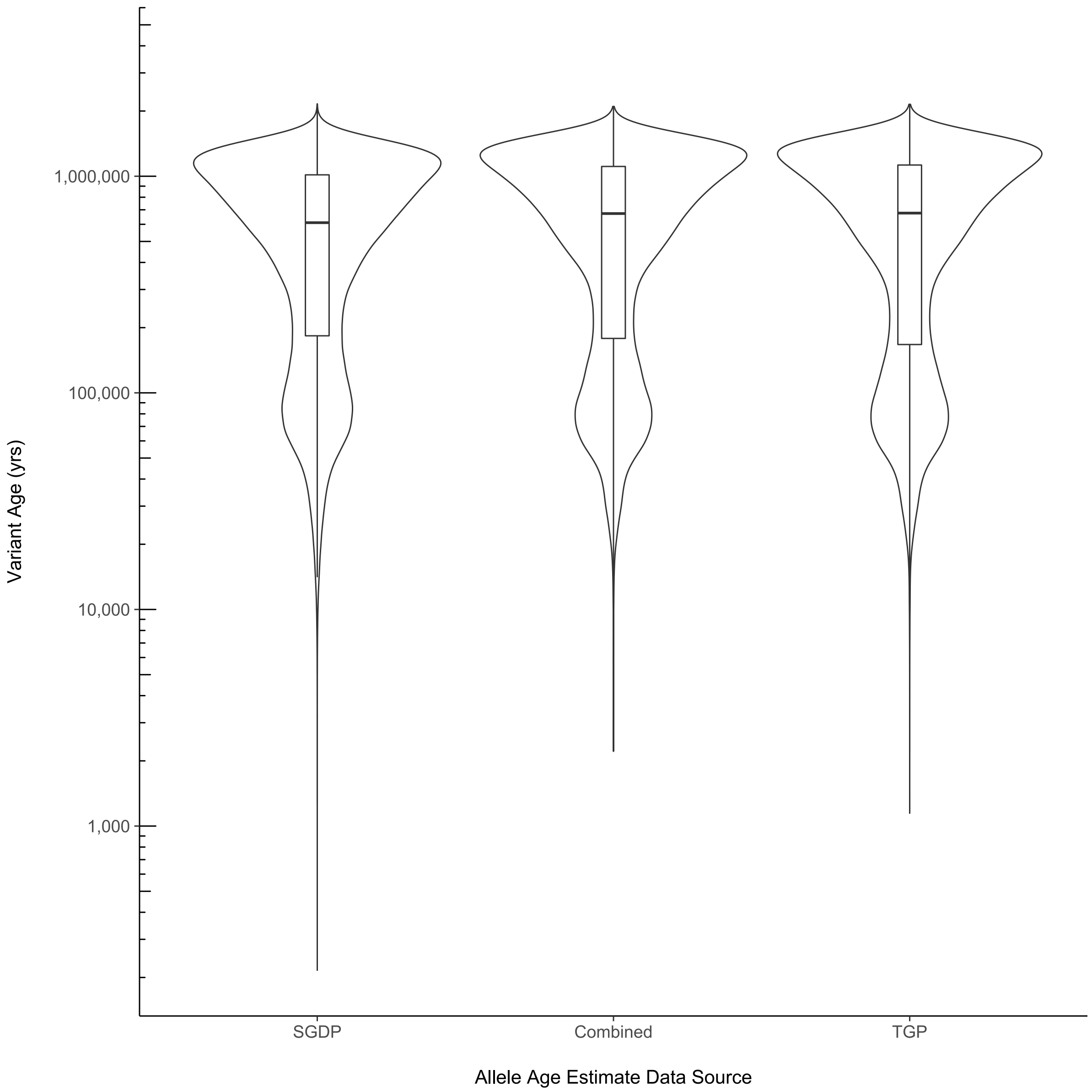


**Fig. S6. AGV age estimate distributions are similar across different data sources.** AGV age estimate distributions for all AGVs present in AVA with estimates from all three data sources: SGDP—Simons Genome Diversity Project, TGP—Thousand Genomes Project, and Combined—both SGDP and TGP (N = 637,502). Ages are the mode of the joint clock estimation displayed in years. Distributions are visualized as violin plots and boxplots where the boxes indicate the median and IQR, with the upper whiskers extending to the largest value $\leq$ 1.5 x IQR from the 75th percentile and the lower whisksers extending to the smallest value $\leq$ 1.5 x IQR from the 25th percentile. Outliers are not shown in the boxplots.

#### Supplemental Tables

| Population | MAF Class | $R^2 \geq 0.2$ | $R^2 \geq 0.5$ | $R^2 \geq 0.6$ | $R^2 \geq 0.7$ | $R^2 \geq 0.8$ | $R^2 \geq 0.9$ | $R^2 \geq 0.92$ | $R^2 \geq 0.94$ | $R^2 \geq 0.96$ | $R^2 \geq 0.98$ | $R^2 \geq 1$ | Sum |
|---|---|---|---|---|---|---|---|---|---|---|---|---|---|
| AFR | UR | 1135209 | 643837 | 372274 | 349570 | 349570 | 349570 | 349570 | 349570 | 349570 | 349570 | 349570 | 32165070 |
| AFR | R | 1549944 | 628429 | 532138 | 435018 | 356358 | 277326 | 260176 | 247282 | 231699 | 228122 | 228122 | 14457324 |
| AFR | LF | 3102526 | 1290247 | 1048892 | 863875 | 698598 | 522219 | 479430 | 428070 | 368150 | 281545 | 206709 | 7103365 |
| AFR | C | 8033882 | 5985626 | 5357188 | 4710670 | 3991442 | 3116337 | 2895375 | 2640908 | 2325497 | 1851755 | 567216 | 9090580 |
| EAS | UR | 1308976 | 1027423 | 779722 | 779722 | 779722 | 779722 | 779722 | 779722 | 779722 | 779722 | 779722 | 19553587 |
| EAS | R | 1164863 | 829613 | 695518 | 625110 | 581846 | 526434 | 519890 | 507644 | 506172 | 506172 | 506172 | 9164135 |
| EAS | LF | 1214328 | 967195 | 913886 | 856989 | 788120 | 680099 | 644810 | 604087 | 537512 | 452932 | 405850 | 2268582 |
| EAS | C | 5622288 | 5160208 | 4970029 | 4746858 | 4450699 | 3952501 | 3788008 | 3570656 | 3252480 | 2673157 | 1154193 | 5899580 |
| EUR | UR | 5367310 | 2992388 | 2026523 | 1699717 | 1501551 | 1293541 | 1255617 | 1229263 | 1204432 | 1199025 | 1199025 | 140512443 |
| EUR | R | 1343295 | 934561 | 836898 | 734178 | 617535 | 467551 | 427916 | 378168 | 324162 | 240885 | 163188 | 6864507 |
| EUR | LF | 1575852 | 1116159 | 1030738 | 948719 | 858130 | 729136 | 689657 | 638476 | 567938 | 447884 | 90200 | 2794230 |
| EUR | C | 6095540 | 5566854 | 5346420 | 5087203 | 4749915 | 4213108 | 4046284 | 3832572 | 3531149 | 2992624 | 350641 | 6361836 |
| SAS | UR | 0 | 0 | 0 | 0 | 0 | 0 | 0 | 0 | 0 | 0 | 0 | 0 |
| SAS | R | 1861934 | 834491 | 817215 | 777939 | 748897 | 748897 | 748897 | 748897 | 748897 | 748897 | 748897 | 13361455 |
| SAS | LF | 1650499 | 1178142 | 1082707 | 987313 | 877345 | 729425 | 698955 | 664639 | 626569 | 626569 | 626569 | 3125797 |
| SAS | C | 6276413 | 5648315 | 5384482 | 5072427 | 4663117 | 4005005 | 3792823 | 3526223 | 3150674 | 2586781 | 1954247 | 6585278 |

**Table S1. | Many common variants occur in LD with AGVs.** The number of TopLD variants in LD with AGVs at varying LD thresholds, stratified by ancestry group and minor allele frequency class. UR = ultra rare; R = rare; LF = low frequency; C = common.

| Sample | LD Threshold | N Partial Matches | N Complete Matches | N Evaluated | Partial Matches Proportion | Complete Matches Proportion |
|---|---|---|---|---|---|---|
| Loschbour | 0.5 | 28480984 | 27150193 | 28681135 | 0.993022 | 0.946622 |
| Loschbour | 0.6 | 22365711 | 21636605 | 22456534 | 0.995956 | 0.963488 |
| Loschbour | 0.7 | 17836090 | 17466847 | 17872498 | 0.997963 | 0.977303 |
| Loschbour | 0.8 | 13982723 | 13819534 | 13994787 | 0.999138 | 0.987477 |
| Loschbour | 0.9 | 10148529 | 10098515 | 10151464 | 0.999711 | 0.994784 |
| Ust'-Ishim | 0.5 | 31782846 | 28972187 | 32168071 | 0.988025 | 0.900650 |
| Ust'-Ishim | 0.6 | 24957983 | 23220242 | 25166294 | 0.991723 | 0.922672 |
| Ust'-Ishim | 0.7 | 19914922 | 18889511 | 20017786 | 0.994861 | 0.943636 |
| Ust'-Ishim | 0.8 | 15620562 | 15061695 | 15662043 | 0.997351 | 0.961669 |
| Ust'-Ishim | 0.9 | 11329693 | 11096033 | 11342528 | 0.998868 | 0.978268 |

**Table S2. | High-coverage ancient genomes validate the co-occurrence of AGVs and modern variants in recent human history.** The numbers and proportions of AGV-modern variant pairs that were partial or complete matches in two ancient genomes: Loschbour and Ust'-Ishim at varying LD thresholds. We used $R^2$ to measure LD.
